# CellMet: Extracting 3D shape metrics from cells and tissues

**DOI:** 10.1101/2024.10.11.617843

**Authors:** Sophie Theis, Mario A Mendieta-Serrano, Bernardo Chapa-y-Lazo, Juliet Chen, Timothy E Saunders

## Abstract

During development and tissue repair, cells reshape and reconfigure to ensure organs take specific shapes. This process is inherently three-dimensional (3D). Yet, in part due to limitations in imaging and data analysis, cell shape analysis within tissues have been studied as a two-dimensional (2D) approximation, *e*.*g*., the *Drosophila* wing disc. With recent advances in imaging and machine learning, there has been significant progress in our understanding of 3D cell and tissue shape *in vivo*. However, even after gaining 3D segmentation of cells, it remains challenging to extract cell shape metrics beyond volume and surface area for cells within densely packed tissues. In order to extract 3D shape metrics, we have developed CellMet. This user-friendly tool enables extraction of quantitative shape information from 3D cell and tissue segmentation. It is developed for extracting cell scale information from densely packed tissues, such as cell face properties, cell twist, and cell rearrangements. Our method will improve the analysis of 3D cell shape and the understanding of cell organisation within tissues. Our tool is open source, available at https://github.com/TimSaundersLab/CellMet.

## Introduction

Organs have specific shapes and sizes [1], with morphologies that ensure efficient function, such as the highly branched structure of the lungs enabling rapid gas exchange with the blood. Understanding how dense, three-dimensional (3D) organs form remains a major challenge in developmental biology [2, 3, 4, 5]. This is in part due to the diversity of cellular processes that shape an organ, including division, migration, fusion, and extrusion [6, 7, 8, 9, 10]. Understanding cell shape dynamics is not restricted to development. For example, post-birth injuries require wound repair, whereby cells must reconfigure and populate the damage area [11, 12] while undergoing long-rang migration [13]. Finally, diseases such as cancer can induce shape changes in affected cells and tissues [14].

The physical shape, mechanics and functions of a cell are closely interrelated [15]. Further, the cell genetic programme is now known to respond to mechanosensitive inputs; morphogenesis and cell fate can influence each other during development and repair [16, 17]. Such interactions can lead to self-organising pattern formation [18]. Cell morphological changes can be induced by local rearrangements involving acto-myosin modification [19, 6]. By localising to specific surfaces, these processes can generate distinct cell shape changes [20, 21]. However, effects of boundaries (*e*.*g*.,, the extracellular matrix (ECM)) and forces from neighbouring tissues can also impact cell and tissue shape [22, 23, 24]. Therefore, extracting quantitative information about cell and tissue morphology in 3D is an important challenge.

It has recently been shown that curvature can induce cell reshaping within tissues in 3D [25, 26, 27]. Other processes, such as cell division [28, 29, 30] or orientated rearrangements [31] can also lead to significant variation in 3D cell shape. Recent theoretical work has highlighted the diverse array of possible cell shapes, even while maintaining confluency within tissues [32]. Quantifying these multi-scale processes is challenging, as both segmentation and quantitative analysis become increasingly difficult.

With the development of deep learning based pipelines for 3D cell segmentation, such as CellPose [33] or CartoCell [34], the capacity to analyse how cells organise within a tissue in 3D has substantially increased. Current tools to analyse such cell segmentation data have focused on describing cell morphology [35, 36], employing methods such as spherical harmonics [37]. However, when considering how cells pack within dense tissues it is necessary to extract fine-grained information, such as how edges and faces vary in morphology, and the number of cell neighbours (connectivity) at different locations within the tissue. Here, we introduce *CellMet* to address these challenges of extracting quantitative cell shape metrics, which retain the structure of the cell, especially within densely packed tissues. CellMet enables quick and reliable extraction of relevant morphological measures from 3D cell segmentation. We apply CellMet to a range of biological systems to demonstrate its versatility. This information can be powerful for addressing questions related to tissue material properties [38, 39], inferring forces *in vivo* [40] and comparing with predictions from vertex models, for example. [41, 42]. CellMet is written as a free Python package available at https://github.com/TimSaundersLab/CellMet.

## 1. Methods

### Image acquisition

All images were visualised using an Olympus spinning disk confocal system with a confocal scanning W1 unit (Yoko-gawa), except the *Drosophila* mesoderm invagination described in [43]. Either 60x/1.30NA or 40x silicon immersion objectives were used, with a Hamamatsu ORCA-FusionBT C15440 Digital Camera.

#### *Drosophila* mesoderm invagination

Images on *Drosophila* mesoderm invagination are from [43], used with their permission.

*Drosophila* **heart**

A stage 16 *Drosophila* embryo resulting from the cross of lines TSD0019 (w*; UASp*>*CIBN-GFP; Sb/TM3,Ser) and TSD0009 (HandGal4/TM3 (III)) was dechorionated using 50% bleach, attached on the dorsal side to a 35 mm glassbottom imaging dish (FluoroDish FD35, World Precision Instruments) using Heptane glue and covered with Halocarbon oil 700 (H8898, Sigma-Aldrich),

#### Zebrafish myotome

Whole wild-type zebrafish embryos were injected with *lyn-gfp* mRNA for labelling cell membranes. Injected embryos were incubated at 25°C until they were visualised. Embryos were mounted in low melting agarose and were bathed with embryo medium containing 0.003% tricaine.

#### hESCs on micropatterns

Micropatterned colonies were generated according to [44]. Briefly, plain coverslips were incubated with PLL(20)-g[3.5]-PEG(5) and placed on a UV-Ozone activated chrome mask, then exposed to UV for 8 minutes to generate micropatterned coverslips, which were then coated with 10% rh-Laminin-521 (v/v, Thermo Fisher Scientific). Human ESCs (H9 line - Wi-Cell) were passaged using Accutase (Stemcell Technologies), diluted in StemFlex with 10 µM Y-27632, and adjusted to 600,000 cells/mL. After 3 hours, media was replaced with fresh StemFlex medium. 3 days later (∼70 hours), micropat-terned hESC colonies were fixed in 4% paraformaldehyde and stained using anti-human beta-catenin antibody (R&D Systems) and secondary antibodies conjugated with Alexa Fluor 647.

### 3D segmentation with CellPose

Single cell segmentation was performed using the Cellpose 3.0 algorithm [33]. We generated custom models to apply to our data, starting from the *cyto3* model. For each tissue, we then performed manual correction using the Cellpose GUI. Final models were used for prediction of volumetric stacks creating 2D labels on each XY slice. These were then stitched together using the following parameters (diameter=30, flow threshold=0.4, cellprob threshold=0, stitch threshold=0.1). Labels were saved as multi-page tif files for analysis.

### Generating artificial training images

To generate an artificial segmented tissue we used the Voronoi diagram from the Scipy Python package to obtain a 2D cell pavement. This pavement was duplicated on several slices to create a 3D labelled image. Seeding point density was varied to obtain more or less cells as required.

For image size comparison, we generated an image with 500 *×* 500 *×* 150 pixels and with ∼70 cells. Using the scale function in ImageJ, we changed the image size to 250 *×* 250 *×* 150 and 1000 *×* 1000 *×* 150 pixels. This enabled us to compare the same cell topology with different image size.

For cell number comparison, we generated an image with 500 *×* 500 *×* 150 pixels with ∼150 cells. We then removed border cells to reach two new datasets with ∼70 and ∼30 cells, keeping the same average cell size.

### Software design

CellMet is written in Python and has been developed on Gnu/Linux. It has been tested on Windows and Mac operating systems. Code and installation can be found here: https://github.com/TimSaundersLab/CellMet. We provide example Jupyter notebooks to explain CellMet use.

#### Prerequisites

CellMet requires 3D segmented images to perform the analysis. Here, we refer to a segmented image as a labelled image where each pixel value corresponds to a specific identified cell (ID). As CellMet does not correct for mis-segmentation; this need to be done prior to analysis. CellMet works by decomposing the labelled image into single cell images for faster processing. To optimise digital storage, these cell images are stored as NPZ Numpy’s compressed array format. Each cell object can alternatively be exported as .obj or .ply files, which is compatible with other 3D rendering software such as Blender or Paraview (Fig. 1).

**Figure 1.**
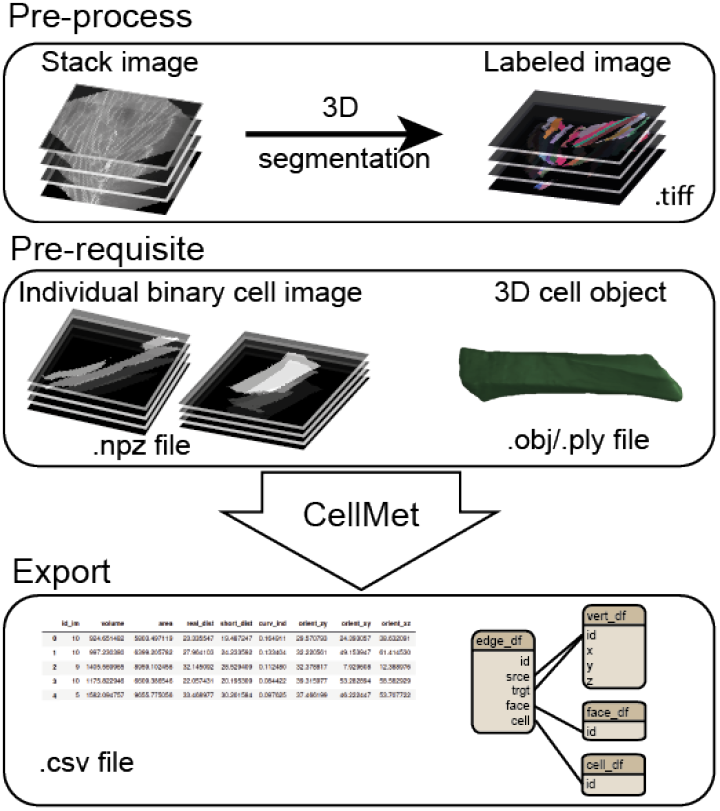
CellMet pipeline. **Preprocess:**Acquisition of a Z-stack. 3D cell segmentation is performed to obtain a 3D labelled image. **Prerequisite** Create single cell files for faster analysis. Possibility to create 3D object file for 3D rendering. **CellMet analysis** Analysis of the shape of cells and neighbour relationship. **Export** Results are stored in .csv files.

#### Decomposing cell, edge and face segmentation to a vertex - like structure

Our approach decomposes the cell into smaller, more manageable structures. We use a half-edge vertex model structure [45] to store data and link them together. The edge table is used to link cell and face tables. This approach enables us to store cell shape information in terms of straight line polygons. To preserve cell topology data such as shape and curvature, additional csv files are saved which contain information about the pixels belonging to each face, edge and cell (Fig. 1, Fig. S1).

To determine cell neighbours, we assume that cells are in close contact with each other. We use the one-cell binary image and dilate by 1 pixel. We then multiply this image by the labelled image. The remaining pixels then belong to the neighbouring cells and their values correspond to the cell ID. Our approach requires cells to be tightly packed; CellMet is not currently easily applicable to tissues with large intercellular spaces.

The second step consists of determining cell edges. At vertices, these edges are shared between three cells. For each cell, all possible combinations of three cells are determined. These three-cell binary images are then multiplied together; if they are not neighbours then the output image is blank. Otherwise, the identified non-zero pixels correspond to the junction between those specific cells (Fig. S2C).

Finally, we determine the surface contact between each pair of neighbours. For each pair, we multiply the binary images together. The non-zero pixels correspond to the interface surface between the cells. Though this approach is simple, it can only determine cell faces if the neighbours are aligned; CellMet does not extract surfaces at the segmentation boundary (Fig. S2B).

#### Cell analysis

Utilising our data tables, we can extract the morphological metrics for a specific cell. To accurately measure the volume and area, images are reshaped into an isotropic pixel size.

Volume and surface area are measured by multiplying the number of pixels on the inside or on the outer surface of the cell by the voxel or pixel size respectively. We use exact Euclidean distance transformations [46] to find the centroid of the cell at each plane from apical to basal (note: our approach does not require cells to be orientated apical to basal). From this, we can also calculate the shortest distance to a surface at each internal point, cell curvature. Orientation of the cell is measured with respect to the three different planes: xy, xz and zy. Cell sphericity is extracted according to:

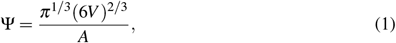

where A and V are the surface area and the volume of the cell respectively.

We do not always require full 3D information about the cell. CellMet can also extract the 2D cell shape at specific z-planes. For each plane we can measure the area, perimeter, orientation, anisotropy, and major axis (Fig. 2).

**Figure 2.**
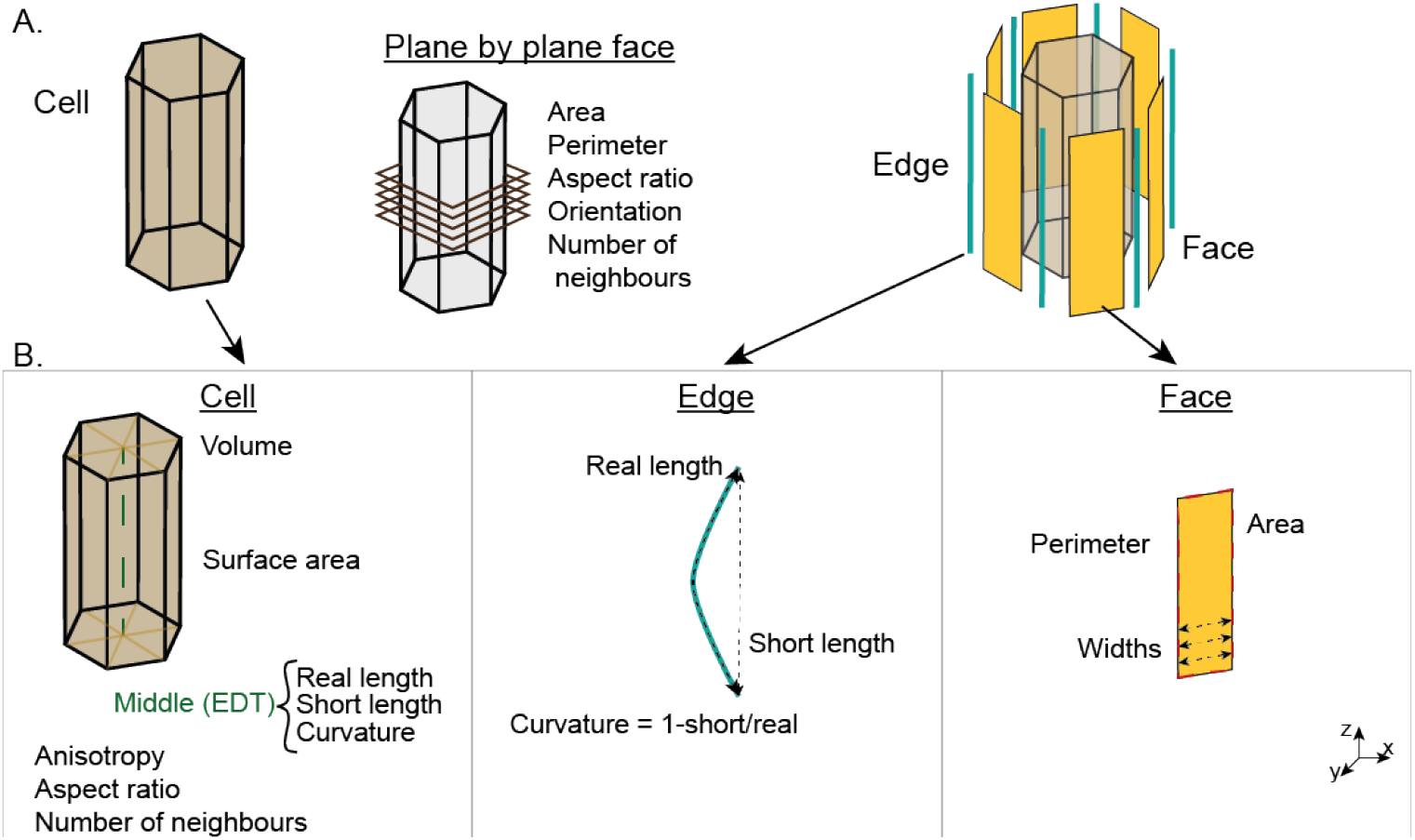
Single cell shape analysis and deconstruction in 2D. A. 3D cell visualisation. B. Metrics description at the level of the cell (left) the edge (middle) and the face (right).

#### Edge and face analysis

For each edge, we can determine the real (*dreal*) and shortest (*dshort*) length (Fig. 2). The edge curvature can be computed as:

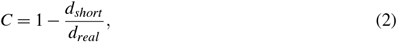

As orientation of edges can be important in a range of biological processes [47], we also extract the length and the angle of the face across each z-plane (Fig. 2).

#### Hardware

For image processing, data analysis and visualisations the workstation used ran Ubuntu 20.04, with 64GB RAM, Nvidia A6000 GPU 48GB and processor Intel Xeon w2235 CPU 3.80GHzx12core.

## 2. Applications

CellMet quickly characterises cell shape in 3D from individual cells that are tightly packed within a tissue. Here, we discuss its applications to a number of example systems and highlight its strengths and weaknesses of our approach.

The shape analysis of 3D cells can be divided into two categories: first, data obtained from each cell individually; second, information about neighbours (Fig. 2). All outputs from CellMet are stored in .csv files (Fig 1).

### *Drosophila* gastrulation

Understanding of tissue morphogenesis -for example, during development or repair - requires knowledge of cell shape within the tissue context. Here, we demonstrate how we use CellMet to extract general cell properties as well as edge positions and surface contacts with neighbouring cells during *Drosophila* gastrulation (Fig. 3). Focusing on the number of neighbours, we observe a significant increment in 3D connectivity during mesoderm invagination (Fig 3). The sharp increase is not observable by considering the apical surface, even though cells undergo a substantial change in size [48]. As we do not consider boundaries, the neighbour number for cells at boundaries is reduced (Fig. 3D).

**Figure 3.**
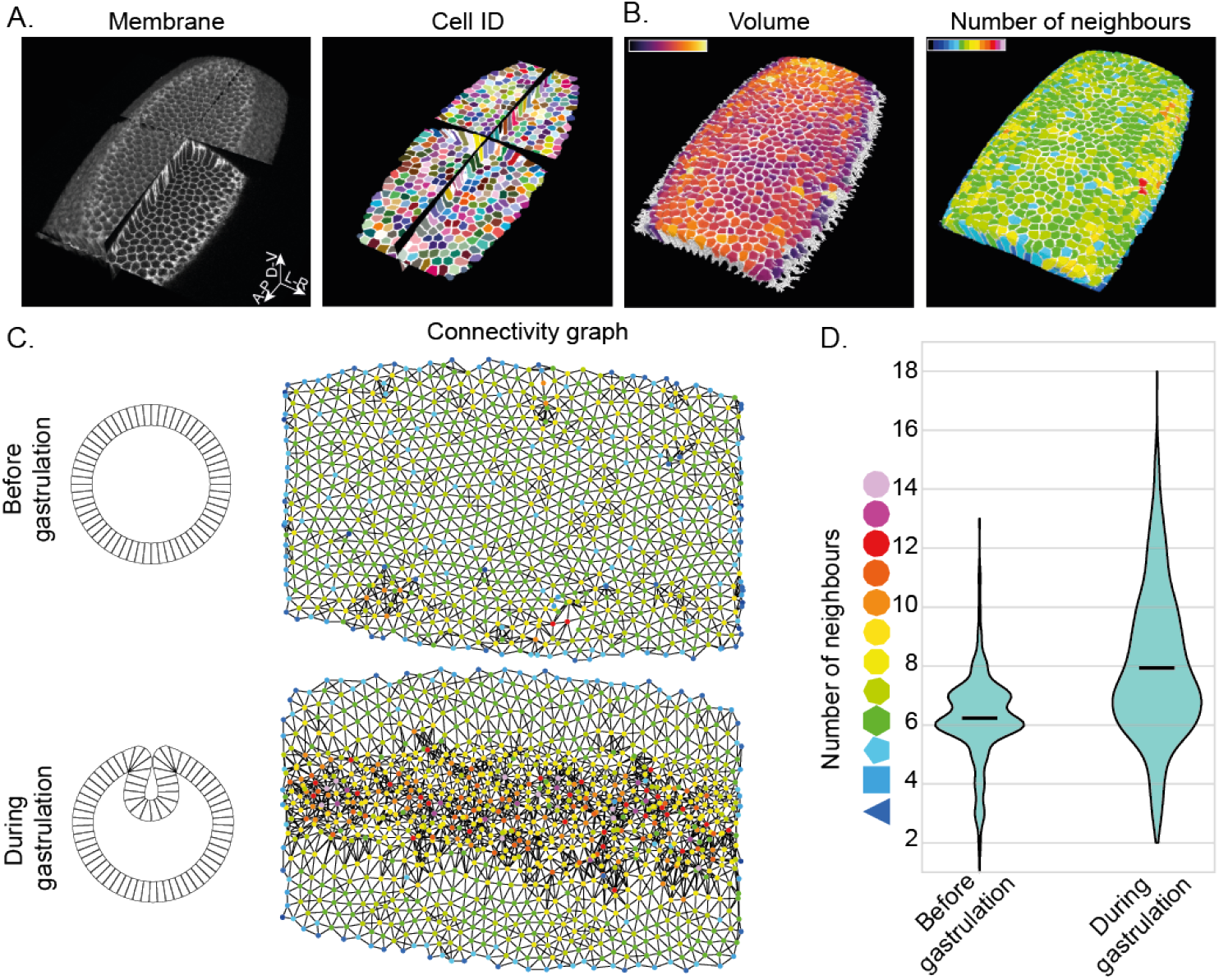
Application to drosophila gastrulation. A. Multi orthoview of PH-mCherry embryo during mesoderm invagination from Gracia [43] (left) and cell identity (right). B. Colour coded cells show volume (left) and neighbour number (right) in the embryo. C. Scheme showing tissue invagination (left) and the corresponding connectivity graph from A. (right) before (top) and during (bottom) gastrulation. Node color represent the number of neighbours and edge color represent the link between two cells. D. Violin plot of the number of neighbours from C. *n* ≃ 900 cells from one embryo.

We can also focus on individual cell shapes. As gastrulation occurs, cells need to rearrange in 3D to stay tightly connected while adjusting for the spatial changes. As a consequence, complex 3D cell shapes, such as scutoids can emerge. Using our single cell analysis, it is straightforward to identify cells that undergo neighbour rearrangements along their apical-basal axis (Fig 4A). We can then extract quantitative information about how cell morphology can alter along its apical-basal axis (Fig 4B).

**Figure 4.**
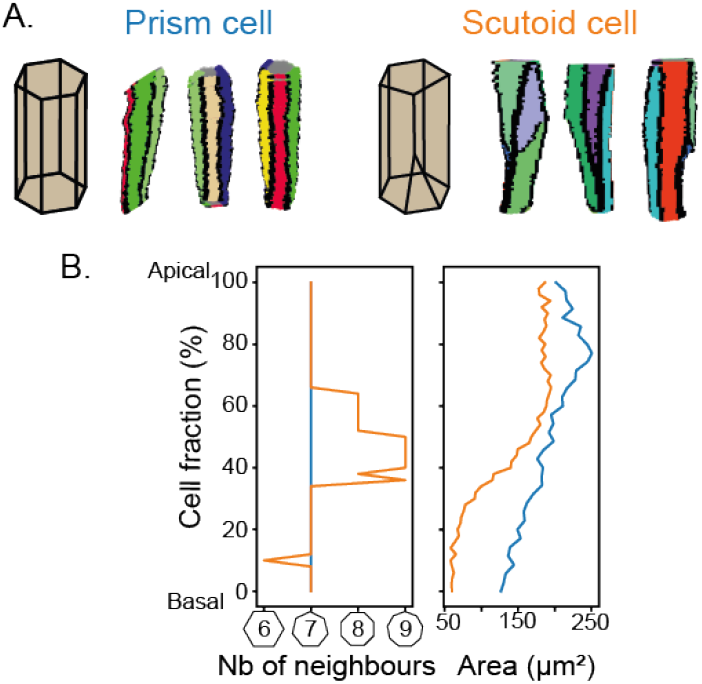
Prism and scutoid cell shape. A. Colour coded face in prism (left) and scutoid (right) cells. Edges are in black. B. Change in number of neighbours and area along cells. Blue and orange represent a prism and scutoid cell respectively.

### *Drosophila* heart

We have extended our analysis to different tissues and model organisms with various organisations. While scutoid geometries have received significant attention, there are other means of 3D cell packing. An example is during *Drosophila* heart closure, where neighbouring cardioblasts must precisely align [49, 50]. We applied CellMet to a stage 16 *Drosophila* embryo heart (Fig 5A). Using our surface deconstruction, we can easily calculate the surface overlap between neighbouring cells. Further, we calculated the connectivity graph (Fig 5A), which reveals information about the precision of cardioblast alignment. This provides a much quicker method to assess cell alignment, compared to human-annotated approaches [51].

**Figure 5.**
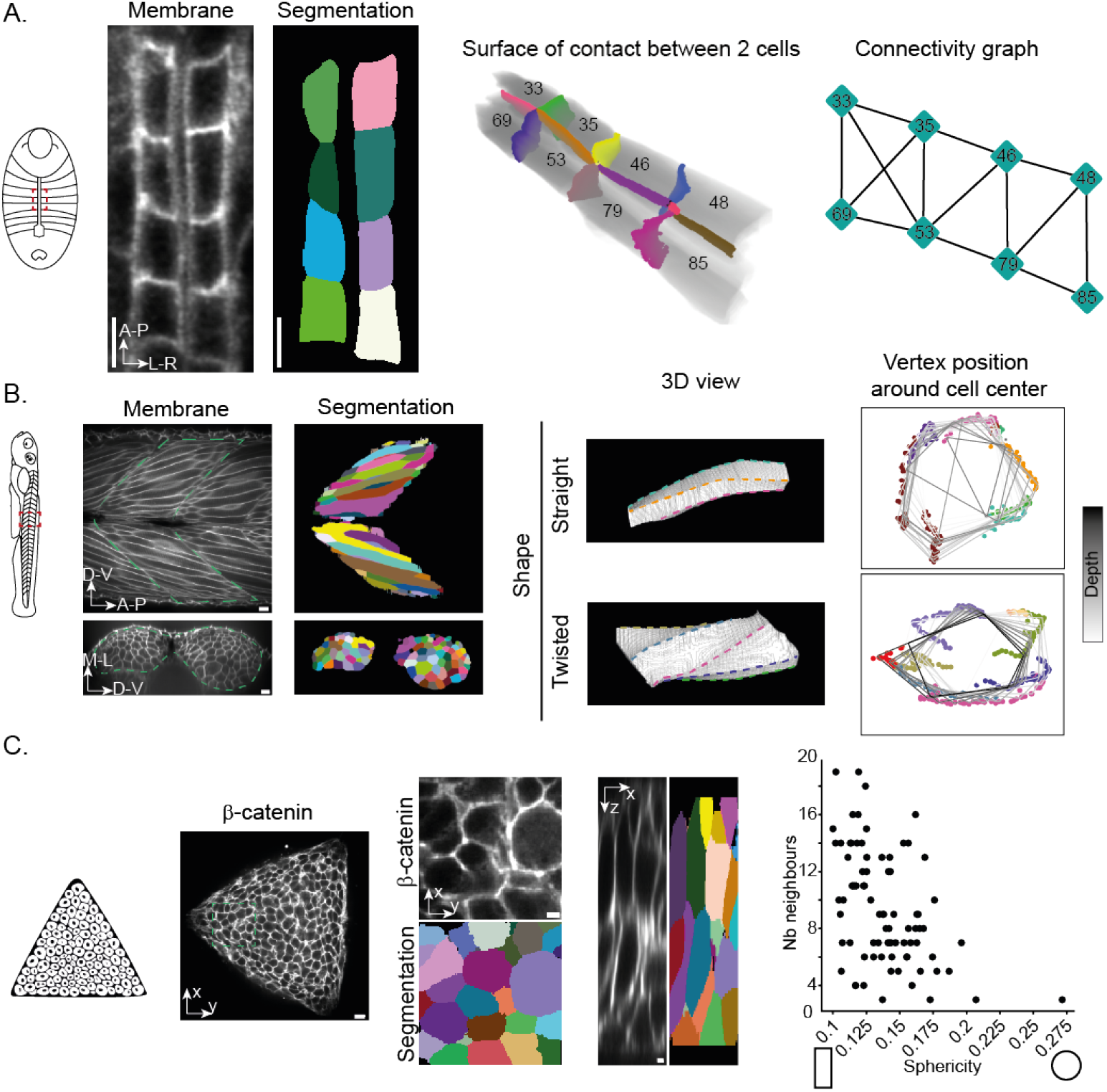
Application of CellMet to different tissues. A. (Left) Schematic of the *Drosophila* embryo from a dorsal view, with anterior up. The red dashed region corresponds to the images to the right. (Centre) Heart cells in a stage 16 *Drosophila* embryo, just as the heart lumen forms with segmentation from CellPose. (Right) Connectivity of the heart cells, demonstrating the cell alignment. Numbers correspond to cell labels ID. B. (Left) schematic of the zebrafish embryo, highlighting the myotome region with red box. (Centre) Myotome in 48hpf zebrafish embryo, with segmentation from CellPose. Green dashed line shows the chevron form of the myotome. Bottom shows cross-sectional view. (Right) 3D visualisation of muscle fibres showing the different cell morphologies. For the twisted cell, we can trace the rotation along the cell long axis. Vertex color are the same as edges showed in 3D. C. (Left) Schematic of organoid culture constrained to triangular domain. (Centre) Organoid culture at 48 hours, with CellPose segmentation. Views shown in x-y and x-z axes. (Right) Comparison of cell sphericity with cell connectivity. Scale bar: A: 5*µm*; B and C: 10*µm*.

### Zebrafish myogenesis

During morphogenesis, zebrafish form muscle segments (myotomes) along their anterior-posterior axis (Fig 5B left). These develop into a characteristic chevron morphology, where muscle cells have to fit closely together [22] in order to synchronously contract to make the fish move.

Using CellMet, we can quickly and precisely highlight contact edges and surfaces along the myotome (Fig 5B right). By analysing cell shape in 3D, we were able to identify a broad range of cell shapes. We observed both straight and twisted cells. In the latter case, cells appear to rotate relative to their neighbours. By tracing the edges of such cells (colour coded dots in (Fig 5B right)), we can quantify their rotation around the cell centre. Thus, CellMet enables analysis of complex cell morphologies that are distinct from 2D behaviours.

### Neuruloids

Organoids have become powerful tools for understanding organ development, including in humans [52]. Yet, detailed analysis of cellular morphodynamics in 3D within organoids remains very limited [53]. Here, we demonstrate that Cell- Met is a powerful tool for extracting relevant morphological information from organoids.

We use an organoid culture, termed neuruloid, which mimics dynamics of stem and progenitor cell populations in the posterior embryo [44]. After 48 hours, the cells become organised in a multilayered epithelium, making it difficult to quantify cell shape (Fig. 5C left). Utilising a trained model in CellPose, we were able to segment the cells in 3D within the multilayered tissue (Fig. 5C centre). Using CellMet, we could then identify the cell interfaces. From this, we built the cell connectivity graph to understand how cells are organised within the tissue. In tandem, we can extract cell shape properties, such as cell sphericity. We see that there is correlation between cell neighbour number and cell shape (Fig 5C right). Overall, Fig. 5 shows that CellMet enables comparison of different cell features, which provides important information for understanding how densely packed tissues are organised.

### Performance testing

To verify the efficiency of CellMet, we generated a synthetic epithelium with a known number of cells (Methods). We created variants of this with different sizes of the same images. Unsurprisingly, larger images increased the time required for analysis. Doubling the image size increased the execution time by a factor of six (Fig 6A). Further, larger images led to memory issues, especially when using parallel execution to quicken the analysis. The execution time also increased with cell number (Fig 6B). The main cause for this slow-down was due to the face-analysis step, as more cells had a large number of neighbours.

**Figure 6.**
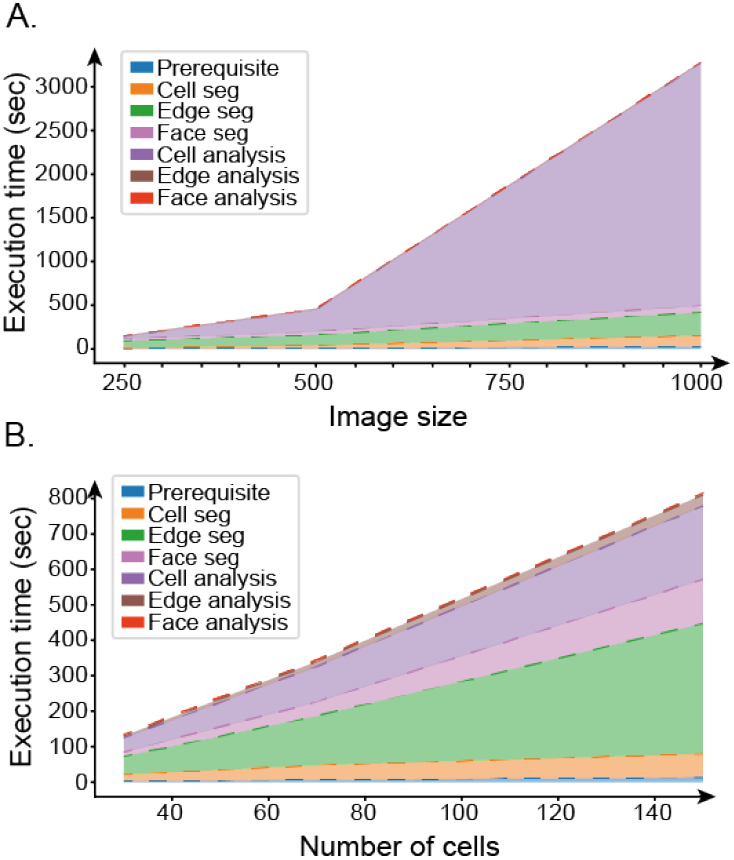
Performance. Execution time on synthetic data using a computer with *n*_*core*_ = 10, and RAM = 64 GB. A. Relationship between the size of the image (*i*.*e*. number of pixels) and the duration of execution. B. Cumulative execution time according to the number of cells in the image. Each subprocess is colour coded (legend), to show the specific time demands.

The accuracy of the analysis is dependant on the quality of the segmentation. CellMet does not correct for mis-segmentation; this needs to be done prior to analysis. Data can be filtered based on given parameters to remove aberrant cells. As stated above, CellMet does not analyse cell faces against the outer boundary. At present, the top and bottom of cells within a monolayer are not considered as faces themselves.

## 3. Discussion

CellMet is a free Python package for extracting quantitative shape information about cells within dense tissues. We provide notebooks with examples showing its application. Cell- Met will work on any segmented data with clear cell identification and sufficient packing.

CellMet provides quick and rigorous analysis of 3D cell properties. It is aimed at understanding cells within densely packed tissues. Therefore, it is not suitable for analysing single cells, *e*.*g*., migrating cells with large protrusions [54]). CellMet is relatively fast, with most analyses completed in under 10 minutes. However, it is not highly optimised, and so slows down with very large images. In this case, data will need to be partitioned to make it more manageable.

In principle, the output from CellMet can be combined with other forms of biological image data. For example, given the edge and face details, signal fluorescence and localisation changes over time can be quantified. This would be relevant for exploring how polarity factors alter their localisation during tissue morphogenesis [55], how morphogens form concentration gradients within dense tissues [56, 57], and how the mechanosensitive protein Piezo1 changes its location and activity levels during organogenesis [58].

The output of CellMet utilises vertex-like approaches to tabulate the results. This means data structure easily permit to link experimental observations with mechanical models of tissue morphogenesis [59]. This information will be important for testing models of how, for example, mechanical stress influences organ shape [60, 61].

The addition of boundary layers will be an important next step. This will enable information from boundary constraints to be more rigorously integrated with cell shape changes. Further, in principle we can include extracellular spaces within the framework (essentially by treating such spaces as specially marked “cells”). The presence of such spaces within even densely packed tissues can play an important role in tissue morphogenesis [38].

In summary, we provide a computational framework for extending our understanding of tissue morphogenesis into three-dimensions. It is aimed at general users; it does not require extensive coding skills.

## Acknowledgements

This work was funded by UKRI Physics of Life Grant (EP/W023865/1) and BBSRC Responsive Mode Grant (BB/W006944/1). We thank James Briscoe, Guillaume Char- ras, Sarah Goodband and Tiago Rito with support in generat- ing organoids and comments on the work. We thank members of the Saunders lab for useful input in the design of CellMet. We thank the Suzanne lab for sharing their data on *Drosophila* embryo gastrulation.

## Author Contributions

S.T. and T.E.S conceived the project. S.T. develop the software. M.A.M-S., J.C. and B.C.Y.L. performed the experiments. S.T. and M.A.M-S performed data analysis. S.T. performed data visualisation. T.E.S. supervised the project. S.T and T.E.S wrote the first draft of the manuscript, with all authors contributing to the final submitted version.

## Supplementary Figures

**Figure S1.**
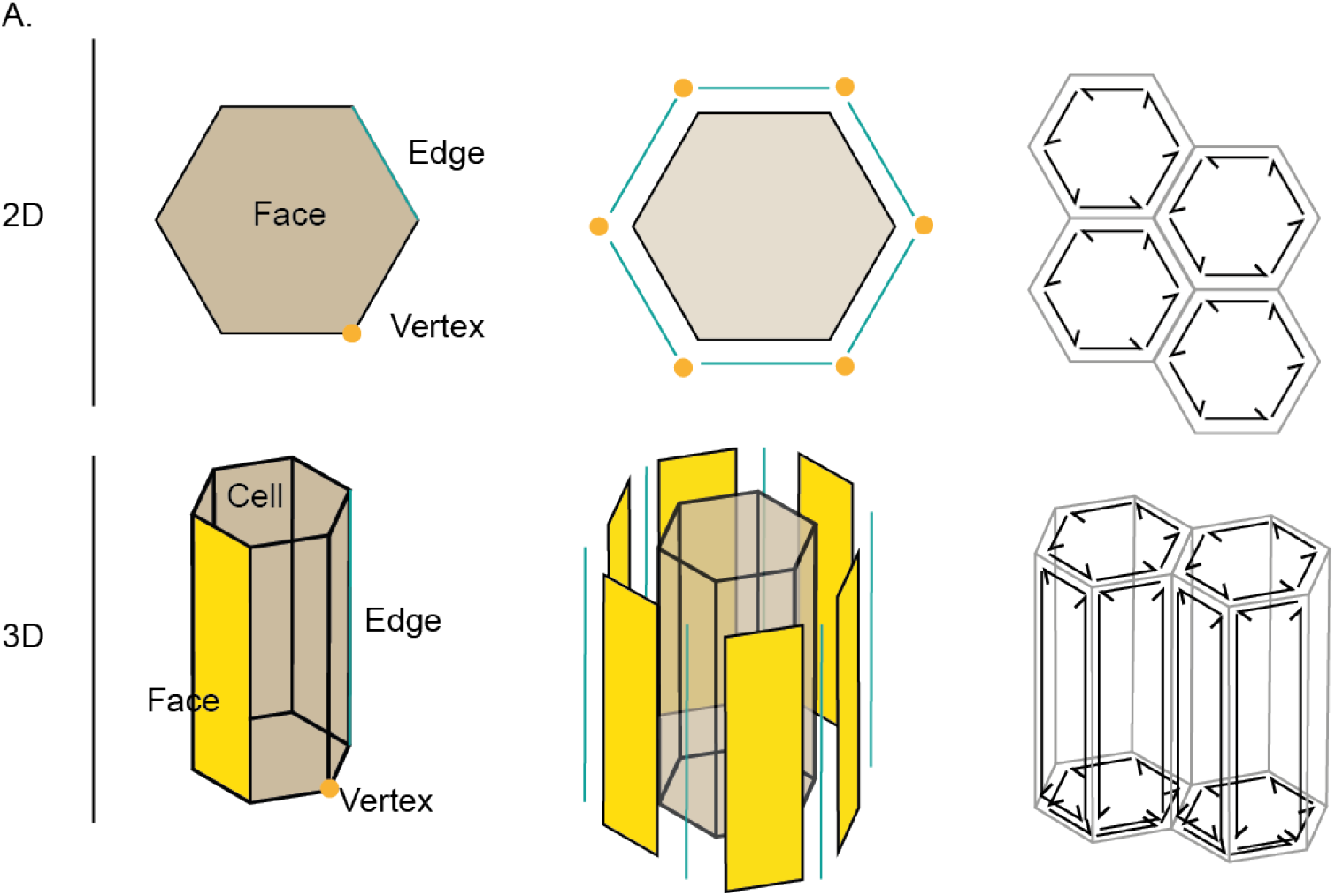
Nomenclature for 2D and 3D cell decomposition. Cell decomposition into faces, edges and vertices in 2D (top) and 3D (bottom).

**Figure S2.**
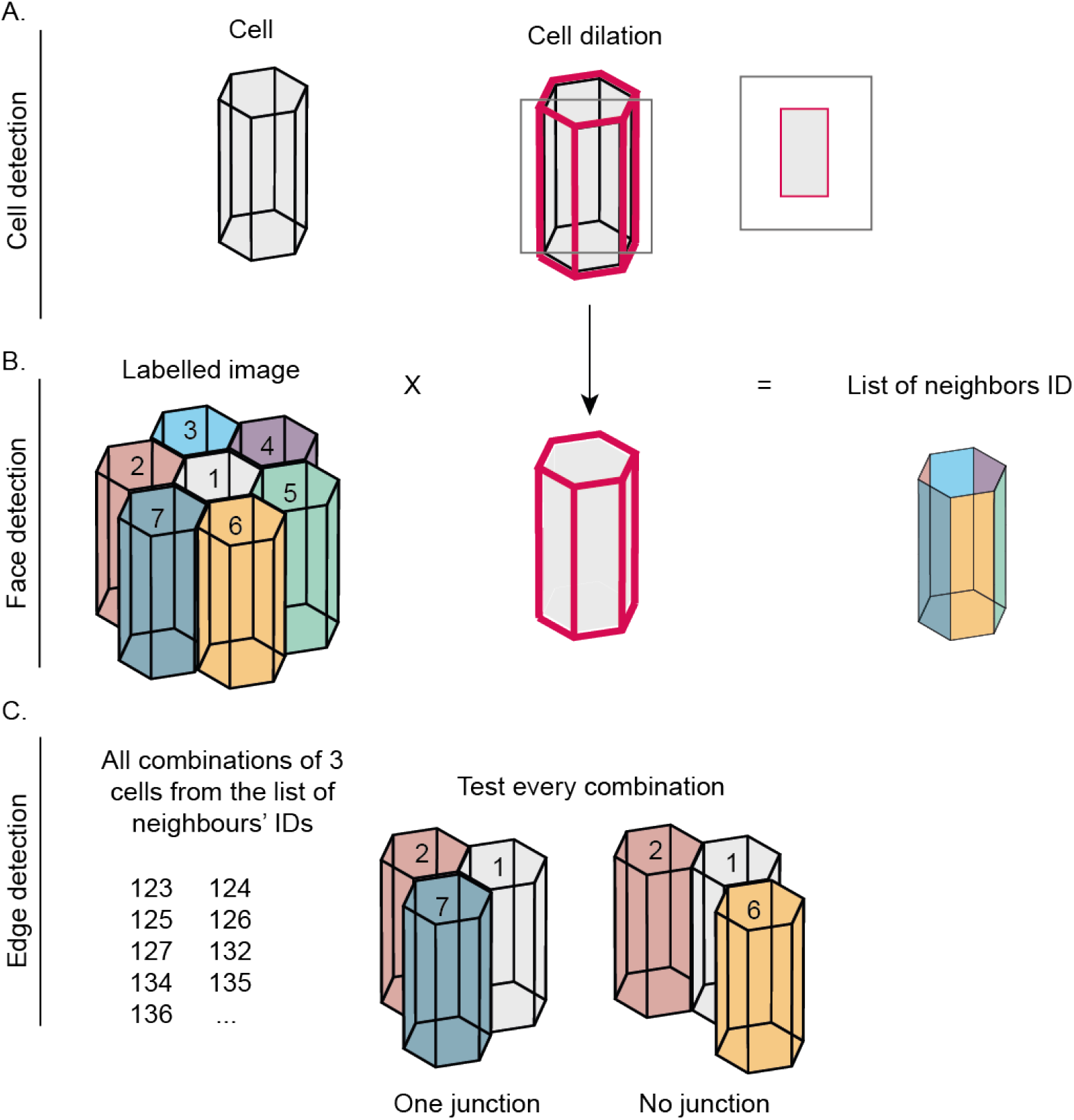
Cell, Edge and Vertex detection. A. Cell detection is based on the segmentation. Cells dilated by one pixel are generated to facilitate face and edge detection. B. Faces are found by multiplying the labelled image by one dilated cell. Non-zero pixel values correspond to the neighbouring cell ID. C. From the list of neighbouring cells, we find all combinations of 3 cells and multiply these 3 dilated cells. Pixels remain when they form a junction, otherwise they are removed to indicate empty space.

